# A Rigid Parallel-Plate Artificial Placenta Oxygenator with a Hemocompatible Blood Flow Path

**DOI:** 10.1101/2022.08.23.505025

**Authors:** David G. Blauvelt, Nicholas C. Higgins, Bianca De, Mark S. Goodin, Nathan Wright, Charles Blaha, Jarrett Moyer, Benjamin W. Chui, Francisco J. Baltazar, Peter Oishi, Shuvo Roy

## Abstract

Extremely preterm infants have poor clinical outcomes due to lung immaturity. An artificial placenta could provide extracorporeal gas exchange, allowing normal lung growth outside of the uterus, thus improving outcomes. However, current devices in development use hollow-fiber membrane oxygenators, which have a high rate of bleeding and clotting complications. Here, we present a novel style of oxygenator composed of a stacked array of rigid and flat silicon semi-permeable membranes. Using computational fluid dynamic (CFD) modeling, we demonstrated favorable hemocompatibility properties, including laminar blood flow, low pressure drop, and minimal cumulative shear stress. We then constructed and tested prototype devices on the benchtop and in an extracorporeal pig model. At 20 mL/min of blood flow, the oxygenators exhibited an average oxygen flux of 0.081 ± 0.020 mL (mean ± standard error) and a pressure drop of 2.25 ± 0.25 mmHg. This study demonstrates the feasibility of a building a stacked flat-plate oxygenator with a blood flow path informed by CFD.

## Introduction

Extreme prematurity, defined as a gestational age of fewer than 28 completed weeks, carries a high burden of morbidity and mortality, in large part due to the underdevelopment of the lungs^1,2^. The transition from the uterine aqueous environment to gas ventilation arrests lung development in an immature stage^3,4^. Furthermore, nearly all extremely preterm infants require positive-pressure mechanical ventilation to ensure adequate gas exchange, which damages the lungs further, a concept known as a ventilator-induced lung injury (VILI)^5^. Ultimately, this leads to inflammation, abnormal lung development, and eventually chronic lung disease, known as bronchopulmonary dysplasia (BPD)^6^.

To mitigate the effects of harmful gas ventilation, there has long been an interest in developing an “artificial placenta” to perform extracorporeal gas-exchange^7^. More recently, several research groups have made advances in development of an artificial placenta circuit using hollow-fiber membrane oxygenators^8–10^. These devices are analogous to extracorporeal life support (ECLS), an advanced therapy for severe respiratory or cardiac failure, in which blood is pumped out of the body and through a hollow-fiber membrane oxygenator before returning to the patient. An extracorporeal artificial placenta can be used to support gas exchange to allow complete lung rest or partial support, enabling lung-protective ventilation, or even non-invasive ventilation. However, one of the primary downsides of hollow-fiber membrane oxygenators is their need for systemic anticoagulation to prevent clot formation. Anticoagulation leads to a high incidence of severe bleeding, including intracranial hemorrhage, which is associated with a mortality of more than 75%^11^. In extremely preterm infants, this risk is significantly augmented due to their underdeveloped and fragile cerebral vasculature, which is prone to rupture and bleeding^12^. For this reason, improving the hemocompatiblity of the oxygenator to allow for minimal or no systemic anticoagulation is critical.

Here, we present a novel style of oxygenator composed of an array of semipermeable silicon membranes, with a blood flow path optimized for hemocompatibility using computational fluid dynamic (CFD) modeling. We show that a composite membrane composed of a semiconductor silicon backbone and a thin elastomeric layer can enable the gas exchange properties of a flexible polymers yet maintain the structural rigidity of semiconductor silicon. We take advantage of the unique rigidity of the flat-plate silicon membranes to create a precisely stacked array of membranes. Using CFD modeling, we demonstrated favorable properties of laminar blood flow, avoidance of stasis and high shear forces, and low pressure drop. We then constructed oxygenator prototypes and tested them in an *in vitro* benchtop mock loop and an *in vivo* extracorporeal pig model to experimentally assess preliminary gas transfer and clotting characteristics.

## Results

### Semiconductor silicon-based membranes enable an oxygenator with a stacked array of membranes

The envisioned artificial placenta circuit consists of an oxygenator connected to the infant via the umbilical circulation (Figure 1). It operates without an external pump, using the pressure differential between the arterial and venous systems to drive flow. Deoxygenated blood from the umbilical artery travels to the device, where oxygen diffuses into the blood and carbon dioxide out of the blood. The oxygenated blood will return to the infant along with medications and nutrition via the umbilical vein. Target specifications for a clinical-scale device are outlined in Table 1.

**Figure 1:**
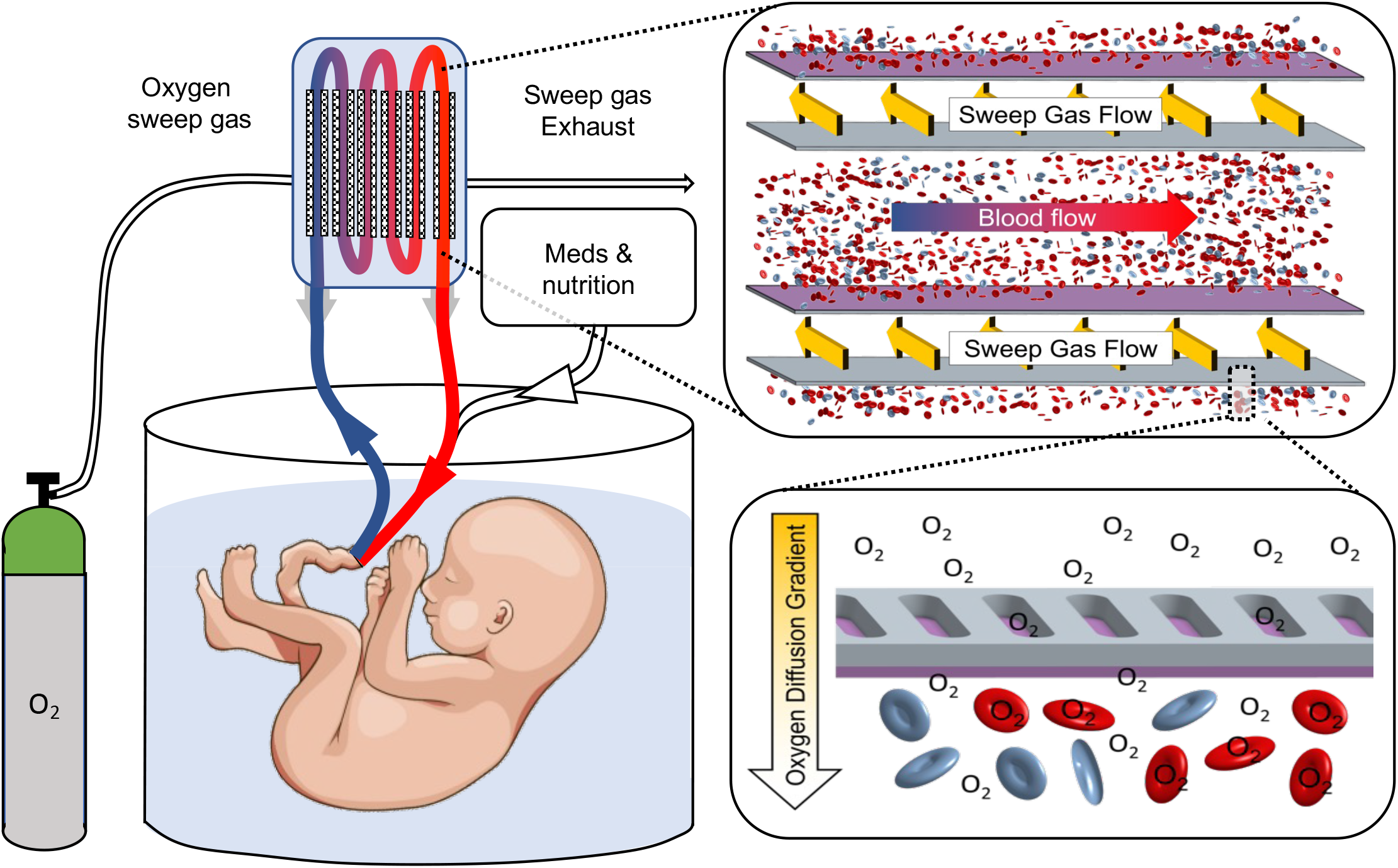
Schematic illustrating the design of a parallel plate oxygenator. The oxygenator is connected to the umbilical circulation and operates without an external pump. Deoxygenated blood is pumped by the infant’s heart out the umbilical arteries to the oxygenator, where gas exchange occurs. The oxygenated blood, along with medications and nutrition, returns to the body via the umbilical vein. The oxygenator itself is composed of a stacked array of alternating blood and sweep gas channels, such that each blood channel is bounded by a sweep gas channel on both the top and bottom. Oxygen from the sweep gas diffuses through the pores of the semiconductor silicon, across the gas-permeable polymer layer (purple), and into the blood.

**Table 1:** Target specifications for an ideal artificial placenta oxygenator

| Specification | Target Value |
| --- | --- |
| Flow rate | 150-200 mL/kg/min |
| Oxygen flux | >6 mL/kg/min |
| Priming volume | <50 mL |
| Pressure drop | 20-25 mmHg |
| Anticoagulation | No anticoagulation |

The oxygenator itself is composed of alternating blood and sweep gas channels, arranged into a stacked array. With this alternating arrangement, each blood channel is bounded by a sweep gas channel on both the top and bottom interfaces. The semipermeable membrane is composed of a pore-containing semiconductor silicon backbone, coated with a gas-permeable elastomeric polymer. Oxygen from the sweep gas channel diffuses down a concentration gradient through the pores, across the gas-permeable elastomeric layer, and into the blood.

### Computational Fluid Dynamic Modeling Shows Promising Hemocompatibility Features

A blood flow path was designed in 3D computer-aided design (CAD), and CFD modeling was performed to evaluate the blood flow path design (Figure 2). The 3D-rendering of the blood flow path and gas exchange membrane windows (Figure 2a) was converted into a mesh with 37.4 million elements. CFD modeling using a blood flow rate of 200 mL/min was then performed to evaluate flow characteristics, pressure drop, and wall shear stress. The blood flow throughout the device was laminar with a maximal Reynold’s number of 271. To evaluate for areas of recirculation or stasis, 100 streamlines were generated, colored by velocity, and followed through the device (Figure 2b). There were no areas of recirculation or stasis, and velocity profiles were well-distributed along the entire blood flow path. CFD was also used to predict pressure drop across the device (Figure 2c,d). Total pressure drop was 5.5 mmHg with largely uniform pressure decreases throughout the flow path. Evaluation of shear stress demonstrated a mean wall shear stress of 0.73 Pa in the channels (Figure 2e). The peak shear stress was 6.3 Pa, well below the number needed to induce red blood cell hemolysis (100 Pa)^13^. Previous literature has demonstrated the role of cumulative shear stress in causing platelet activation and subsequent thrombosis^14–16^. A linear stress accumulation model was created to simulate an average platelet undergoing 3000 passes through the device. A probability density function based on this simulation (Figure 2f) showed that even after 3000 passes, platelet stress accumulation was below the activation limit described by Hellums et al.^13^.

**Figure 2:**
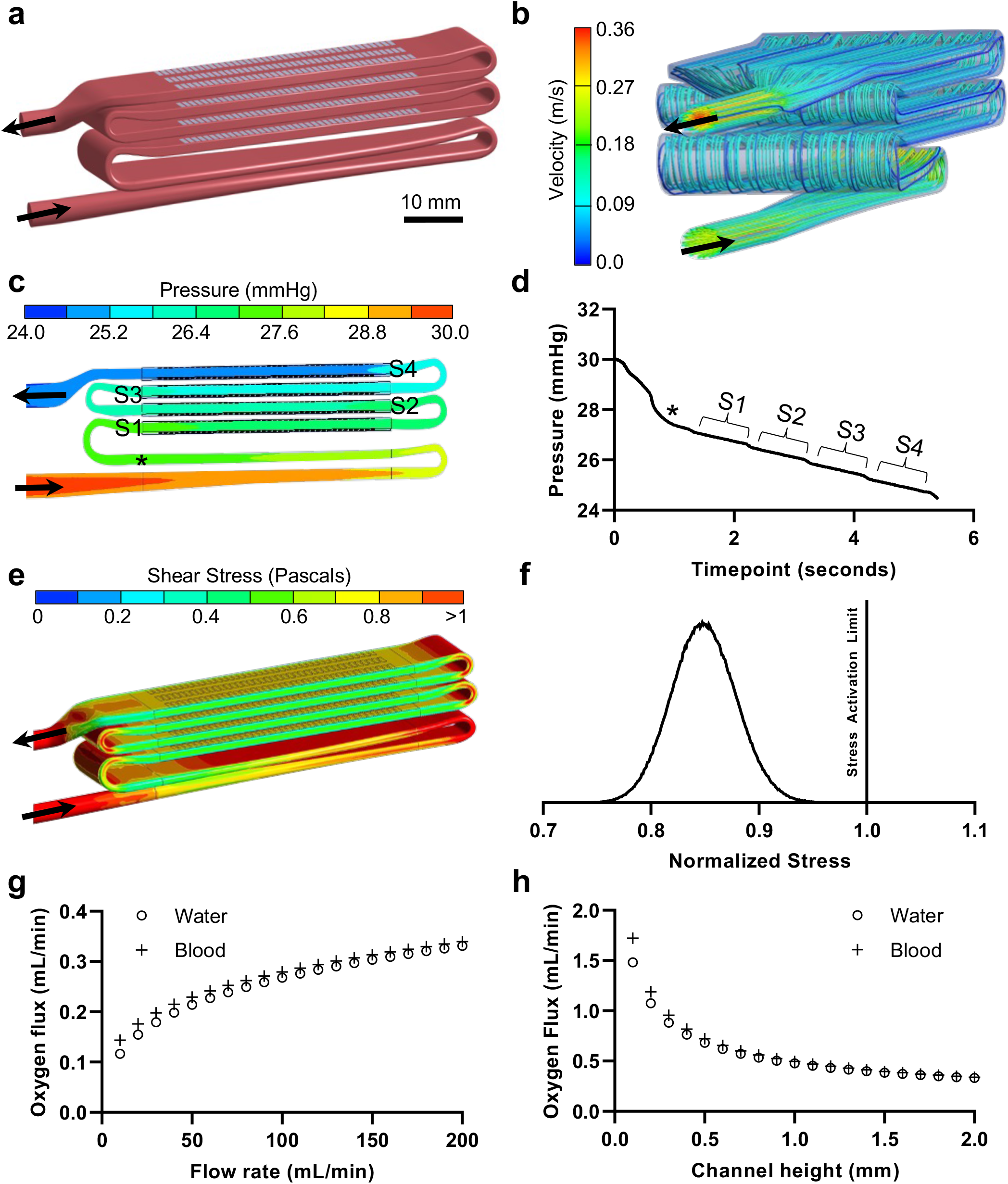
Computational fluid dynamic (CFD) and mass transfer modeling. a) A blood flow path was created using computer aided design (CAD) and used for CFD. Gray rectangles denote areas of active gas exchange. Arrows denote direction of blood flow. b) Streamline analysis using 100 streamlines shows uniform laminar flow throughout the channel, including turnaround points. c) Static pressure variation on the center-line plane demonstrated an area-averaged pressure drop of 5.5 mmHg at a flow rate of 200 mL/min. S1 to S4 refer to the starting points of each segment of gas exchange membranes. d) The largest pressure decrease occurred at the inlet and then was largely uniform through the rest of the device. Asterisk denotes the corresponding marked location in subfigure c. e) Map of the wall shear stress. f) Probability density function of the normalized accumulated shear stress experienced by platelets after 3000 device passes. Cumulative platelet stress remained below the stress activation limit. g) Mass transfer model showing oxygen transfer in water and blood as a function of increasing flow rates h) Oxygen transfer as a function of channel height. Oxygen flux decreases as the channel height increases, with the largest changes occurring at channel heights less than 500 μm.

Oxygen flux into water and blood was predicted using a mass transfer model^17^. Total oxygen flux across the device increased with higher water and blood flow rates (Figure 2g). At 200 mL/min, total oxygen transfer was 0.34 mL/min in the blood model. We also modeled the effect of channel height on oxygen flux (Figure 2h). As the height increased from 100 μm to 500 μm, oxygen flux decreased by 58%, though this efficiency loss tapered off, with another 22% decrease between 500 μm and 2 mm.

### Membranes were fabricated using etched semiconductor silicon and a thin elastomeric coating

The composite membrane consisted of a rigid silicon backbone bonded to a thin layer of gas-permeable polydimethylsiloxane (PDMS). The membranes were fabricated in 3 phases, illustrated in Figure 3. In the first phase (Figure 3a), pores were etched into a silicon on insulator (SOI) substrate to form the rigid backbone. On the top (“pore”) side, photoresist was spin-coated and patterned with micropores of 10 μm wide by 50 μm long, separated by 10 μm on all sides. Vertical trenches were then created by deep reactive ion etching (DRIE) of the pore side silicon down to the buried oxide layer (100 μm depth). The process was repeated on the back (“window”) side except with larger windows of 1 mm wide by 6 mm long by 299 μm deep. The membranes were then wet-etched using hydrofluoric acid, which dissolved the 1 μm buried oxide layer, thereby connecting the pores to the windows. In the second phase (Figure 3b), the thin layer of gas-permeable PDMS was prepared on a multilayered sacrificial substrate. A thick “handle” layer of PDMS was spin coated onto a silicon wafer, followed by a water-soluble polyvinyl alcohol (PVA) layer. Finally, a 5 μm PDMS layer was spin coated on top of the PVA. In the final phase (Figure 3c), the thin PDMS layer was bonded to the silicon backbone using oxygen plasma. The PVA was then dissolved in water, releasing the sacrificial handle layer of PDMS and leaving the completed composite membrane. The composite membranes used for this study (Figure 3d) had overall dimensions of 32 mm wide by 65 mm by 400 μm thick with 7.2 cm^2^ of surface area for gas exchange. Scanning electron microscopy (SEM) demonstrated excellent uniformity of both the pores and PDMS thickness (Figure 3e-g).

**Figure 3:**
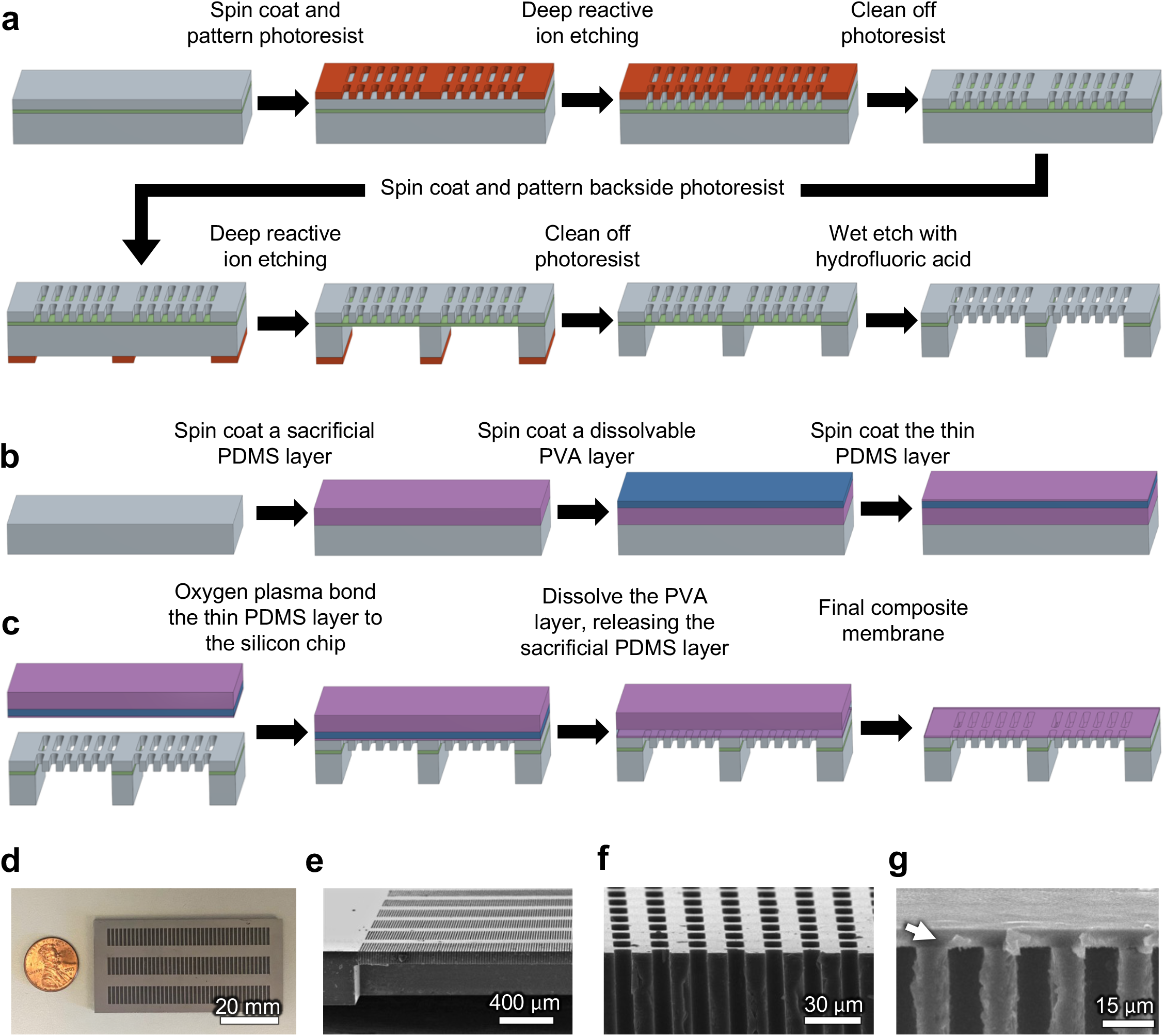
Composite silicone membrane fabrication. a) Silicon on insulator (SOI) is etched using a photoresist pattern and deep reactive ion etching (DRIE). Microscopic pores are etched on the top (“pore”) side and large windows are etched on the bottom (“window”) side. The buried oxide insulator layer is dissolved using hydrofluoric acid, connecting the pores to the large windows. b) The thin (5 μm) polydimethylsiloxane (PDMS) polymer layer is spin coated on top of a sacrificial PDMS handle layer, separated by a layer of polyvinyl alcohol (PVA). c) The final composite membrane is produced by oxygen plasma bonding the thin PDMS layer to the pore side of the etched SOI component. The sacrificial layers are then released by dissolving the PVA layer in water. d) En face view of the composite membrane. e-g) Electron micrographs showing (e) an oblique view of the pores and large windows, (f) a cross-sectional view of a cut through uncoated pores, and (g) a cross-sectional view of a cut through pores bonded to a 5 μm layer of PDMS (arrow).

### The oxygenator was constructed using a stack of silicon membranes and a 3D-printed housing

The oxygenator was composed of a stack of silicon membranes separated by PDMS gaskets and a 3D-printed housing to guide the blood flow through the stack (Figure 4). Although our modeling suggested that a shorter channel height would result in improved flux, fabricating a device with a channel height less than 500 μm was challenged by limitations in the resolution of 3D printing. Therefore, for this initial design prototype, to ensure reproducible device construction, we chose to build and test a device with a 2 mm channel height. The stack was formed by bonding either a 2 mm thick “blood channel” gasket or a 700 μm thick “sweep gas channel” gasket to a silicon membrane (Figure 4a). The gaskets sealed off the edges of the channels, defined the channel heights, and helped provide mechanical support for the stack. To create the completed stack (Figure 4b), 8 silicon membranes and 2 solid silicon end-pieces were bonded together, ultimately forming 4 blood channels, each bounded on both sides by a sweep gas channel. The stack was then slid into the 3D printed housing using thin rails in the housing to guide the alignment of the channels. The seams of the stack were sealed with epoxy to prevent leaking. Sweep gas headers were added to interface with side openings in the sweep gas channels. Finally, side walls and a top plate were screwed into place to add mechanical support, forming the completed oxygenator (Figure 4c).

**Figure 4:**
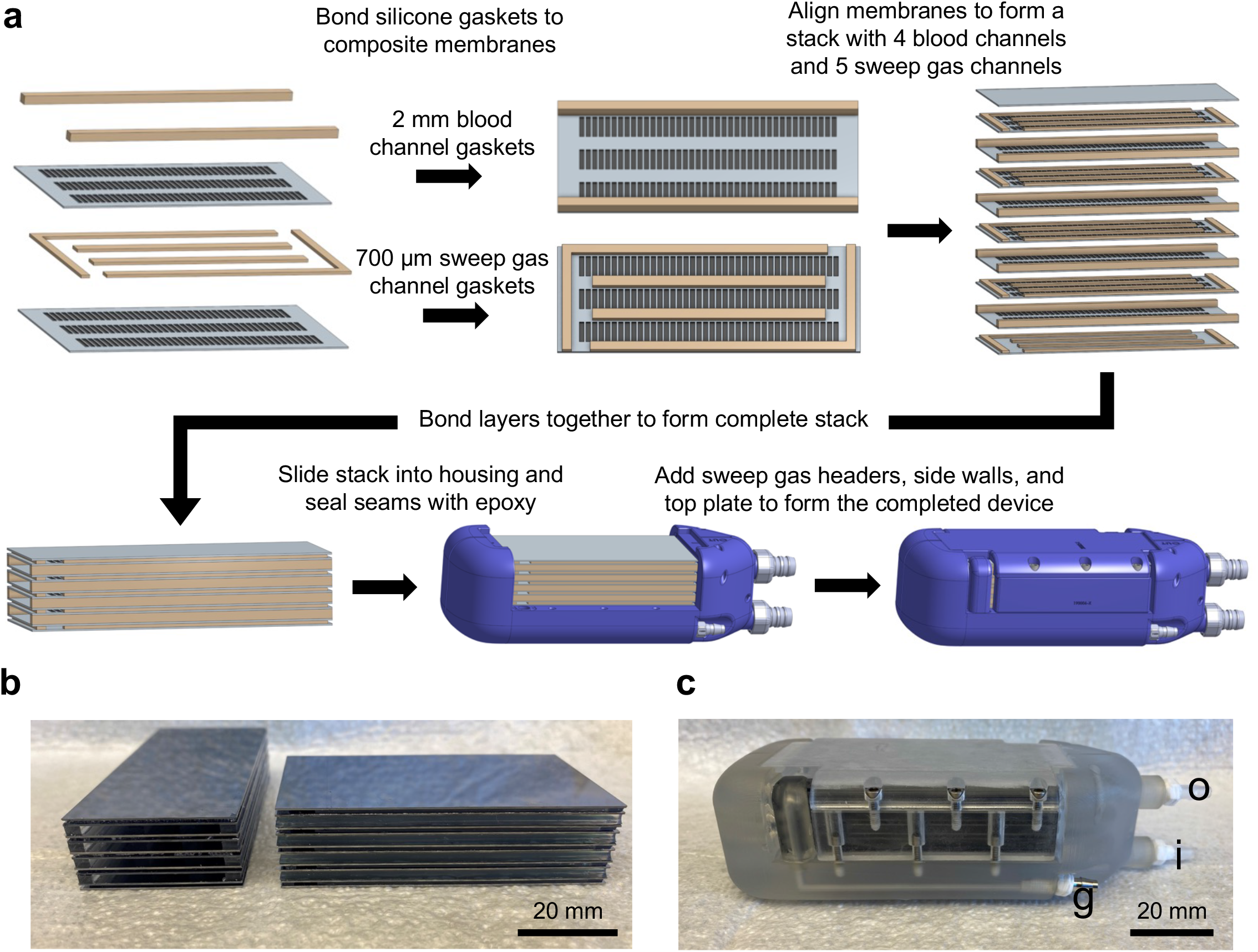
Oxygenator device assembly. a) The membrane stack is formed by bonding silicone gasket spacers, which form the blood and sweep gas channels. The completed stack is composed of 8 total membranes bounded by 2 solid silicon end pieces, which forms 4 blood channels and 5 sweep gas channels. The stack is placed into the housing along guide rails and sealed with medical-grade epoxy. Sweep gas headers are added to direct the sweep gas to the correct channels. Side and top plates provide mechanical support. b) Images of the completed stacks, showing the alternating blood and sweep gas channels. c) Completed device after sealing the stack into the 3D printed housing showing the blood inlet (i), outlet (o), and sweep gas (g) connectors.

### Benchtop and In Vivo Testing Demonstrates Flow Correlated Oxygen Transfer and Low Pressure Drop

Four oxygenators were assembled for benchtop and in vivo testing (Figure 5). In the benchtop testing setup, pressure drop was 0.78 ± 0.15 mmHg (mean ± standard error [SE], n=4 devices), 1.9 ± 0.12, and 3.6 ± 0.21 at 10, 20, and 40 mL/min flow rate (Figure 5c). The partial pressure of dissolved oxygen (mmHg) increased by 173.7 ± 7.2 (mean ± SE, p<0.001 [paired t-test], n=4 devices, 6 measurements per device per flow rate), 122.8 ± 5.4 (p<0.001), and 81.1 ± 3.9 (p<0.001) for 10, 20, and 40 mL/min, respectively (Figure 5d). Oxygen flux was 0.066 ± 0.003 (mean ± SE, n=4 devices, 6 measurements per device per flow rate), 0.094 ± 0.004, and 0.124 ± 0.006 at 10, 20, and 40 mL/min, respectively (Figure 5e). For these experiments, a sweep gas pressure of 0.5-1 PSI was used.

**Figure 5:**
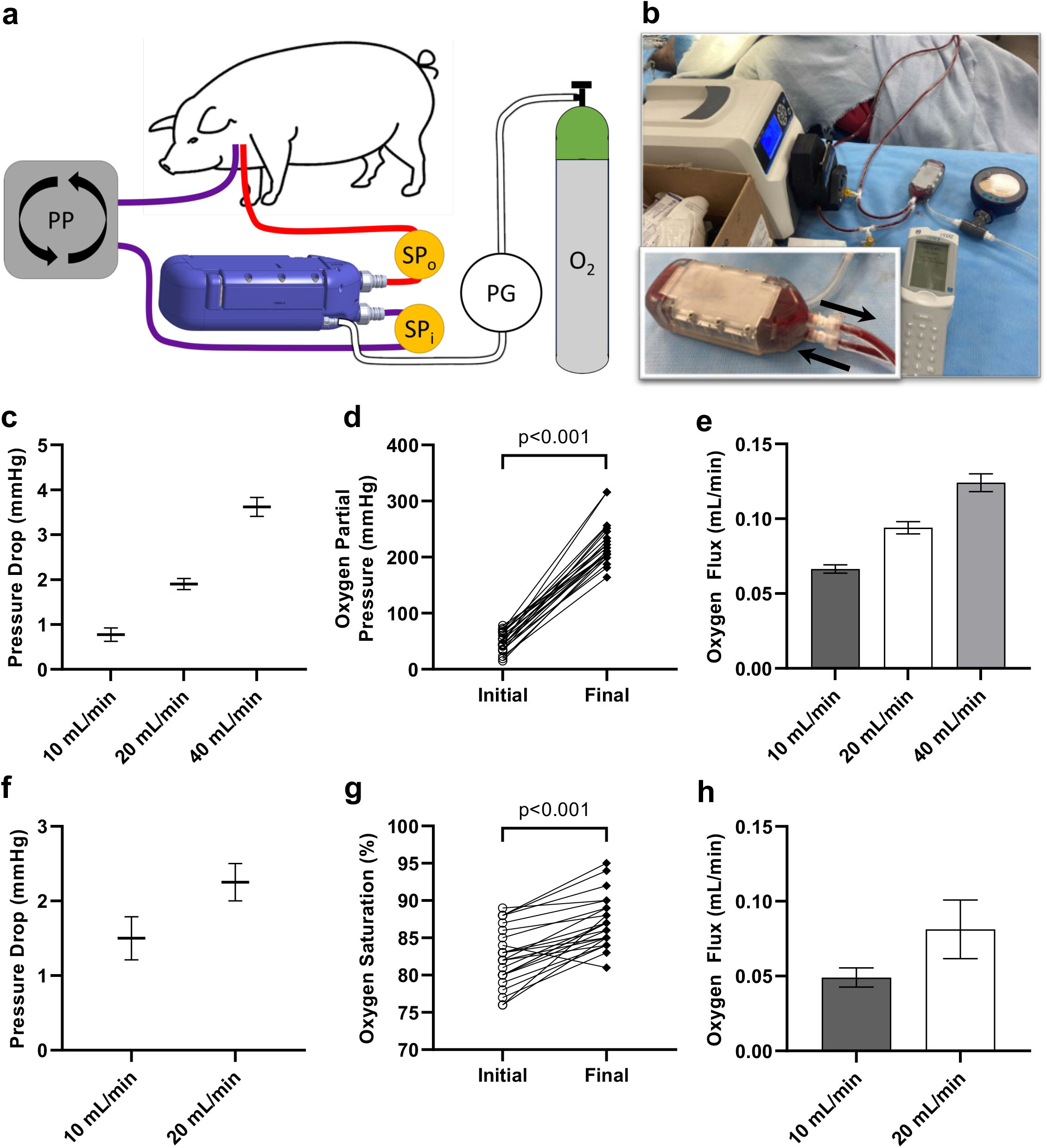
Device testing results. a) Schematic of the device testing in an extracorporeal pig model. Deoxygenated venous blood is pumped to the device via a peristaltic pump (PP). Oxygen sweep gas is connected to the sweep gas port, and the pressure is monitored with a pressure gauge (PG). Pressure measurements and samples for blood gas analysis are taken from the inlet and outlet sampling ports (SP_i_ and SP_o_). b) Image of the device undergoing testing in an extracorporeal pig study. c-e) Device testing in a benchtop water circuit. c) Pressure drop measurements at 10, 20, and 40 mL/min of water flow. d) Partial pressure of dissolved oxygen in water at a water flow rate of 10 mL/min. e) Oxygen flux as a function of water flow rate. f-h) Device testing in an extracorporeal pig model. f) Pressure drop measurements at 10 and 20 mL/min of blood flow. g) Oxygen saturation of hemoglobin at a blood flow rate of 10 mL/min. h) Oxygen flux as a function of blood flow rate. For all experiments, n=4 devices, 6 measurements per device per condition. Data is presented as mean ± standard error. P-values calculated from a 2-tailed paired t-test.

Following benchtop testing, the devices were evaluated in an extracorporeal pig model. A double-lumen venous catheter was placed in the external jugular vein of an anesthetized pig, and blood was pumped through the device using a peristaltic pump. Pressure drop was 1.50 ± 0.29 mmHg (mean ± SE, n=4 devices) and 2.25 ± 0.25 mmHg for 10 and 20 mL/min of blood flow, respectively (Figure 5f). Oxygen saturation (%) increased by 4.75 ± 0.34 (mean ± SE, p<0.001 [paired t-test], n=4 devices, 6 measurements per device per flow rate) and 4.04 ± 1.34 (p<0.001) for 10 and 20 mL/min, respectively (Figure 5g). Oxygen flux *in vivo* closely matched the benchtop measurements (Figure 5h). Mean flux (mL/min) was 0.049 ± 0.006 at 10 mL/min blood flow and 0.081 ± 0.020 at 20 mL/min (mean ± SE, n=4 devices, 6 measurements per device per flow rate). At the end of the 180-minute study, the devices were gently flushed with normal saline at a rate of 10 mL/min to evaluate the device patency. No gross clots were observed in any of the devices.

## Discussion

In this study, we designed and built an oxygenator consisting of a stacked array of silicon membranes for future artificial placenta applications. Previous work in our lab demonstrated that composite silicon membranes could be used to perform gas exchange^17,18^. We built upon this previous proof-of-concept work to create a stacked membrane device featuring a hemocompatible blood flow path. Using CFD, we investigated the blood flow path design to evaluate parameters known to influence hemocompatibility. In addition, we studied the performance of the oxygenator on the benchtop and *in vivo*.

The unique combination of a pore-containing semiconductor silicon backbone with a thin elastomeric gas-permeable layer draws numerous benefits from its mechanical rigidity. Its structural integrity enables the silicon membrane to be easily stacked into a modular structure. Furthermore, the mechanical support offered by the silicon backbone allows the PDMS gas exchange layer to be only 5 μm thick, which offers very little resistance to gas transfer. Finally, the rigid plates enable the use of a higher pressure of oxygen sweep gas, which can potentially further improve gas exchange efficiency.

In addition, the silicon membrane oxygenator incorporates features known to improve hemocompatibility, including a CFD-optimized blood flow path and wide blood channel. Using CFD simulations, we designed a hemocompatible flow path design throughout the entire device, including the inlet, outlet, and flow distribution network. This is critical, as prior studies suggest that these are the components of the oxygenator that are most at risk for flow disruption and thrombosis^19^. CFD modeling demonstrated uniform laminar flow, no recirculation zones, and a platelet stress accumulation below the level needed to activate platelets.

Another important design characteristic is creating a device featuring blood channels with a large cross-sectional width-to-height aspect ratio. Devices featuring conventional square microchannels are prone to clotting due to a high surface-area-to-volume ratio, which allows co-localization of activated clotting factors and platelets to form a stable clot. In addition, microchannels become easily occluded, which alters blood flow dynamics and promotes clot propagation. Therefore, it is necessary to consider strategies to decrease the surface-area-to-volume ratio of the blood channel without sacrificing other important design characteristics. Increasing the height of the channel (i.e. the distance between the membranes) decreases the channel surface-area-to-volume ratio but also has a negative effect on gas-transfer efficiency. Widening the channel for a given channel height decreases the surface-area-to-volume ratio without affecting gas transfer, so a wide channel is a desirable feature. Due to the mechanical rigidity of the semiconductor silicon backbone, this oxygenator was able to feature a very wide channel of 27 mm and a width-to-height aspect ratio of 13.5.

The combination of an efficient semipermeable membrane, a CFD-optimized blood flow path, and blood channels with a high width-to-height ratio have the promise to enable a hemocompatible oxygenator. Although these devices need to be further optimized by incorporating hemocompatible materials and surface coatings, it is important to note that all oxygenators remained clot-free even without systemic anticoagulation. A single dose of heparin (100 units/kg) was given at the time of vessel cannulation and circuit priming, but due to the very short half-life of heparin *in vivo*, the majority of the study (approximately 180 minutes per device) was conducted without therapeutic anticoagulation. By comparison, other PDMS-based oxygenator devices typically require very high-dose anticoagulation (e.g. repeated doses of 400 units/kg of heparin) to avoid device thrombosis ^20,21^.

In the past decade, flexible microfluidic oxygenators have emerged as a potential alternative to hollow-fiber membrane oxygenators. Microfluidic oxygenators, with their capillary-sized blood channels offer significant benefits in gas-exchange efficiency^22^. However, existing microfluidic oxygenators are challenged by many of the issues that the silicon membrane oxygenator is designed to overcome. Flexible microfluidic oxygenators have been difficult to scale up due to the risk of microchannel collapse when stacking multiple layers into a compact device^22,23^. Along with this, microfluidic oxygenators have featured thicker gas exchange layers, typically around 30 to 70 μm, to improve mechanical robustness, with a tradeoff of decreased efficiency^23^.

Furthermore, flexible microfluidic devices have yet to make significant progress in the most clinically relevant parameter for an artificial placenta: hemocompatibility. Although attempts have been made to improve hemocompatibility through the use of biomimetic flow paths^24–27^, microfluidic oxygenators have empirically required very high doses of anticoagulation (typically higher than hollow-fiber membrane oxygenators) to prevent clotting^23^. This finding may be due in part to the combination of their high surface-area-to-volume ratio and non-ideal flow distribution networks. Most microfluidic devices include narrow channel widths on the order of a few hundred microns^25,27^ – two orders of magnitude smaller than the silicon membrane oxygenator – resulting in a high surface-area-to-volume. In addition, although biomimetic flow paths can be integrated into individual layers of a microfluidic device, it has been challenging to integrate hemocompatible designs into a 3-Dimensional flow distribution network designed to distribute blood to the individual layers of a multilayer device. The flow disruptions at the interfaces between the flow distribution network and gas exchange layers create niduses for clot formation and propagation.

A comparison of our device to hollow-fiber membrane oxygenators and flexible microfluidics is shown in Table 2. Our device performs comparatively well in terms of device resistance and anticoagulation need. However, improvements are needed in overall oxygen transfer to achieve a clinical-scale device. Our modeling work suggested that reduction of channel height can improve the diffusion of oxygen from the boundary layer to the bulk flow, thereby increasing overall oxygen transfer. Future designs will focus on finding engineering solutions to reduce the channel height to the sub-millimeter range. Another limitation of our study is the lack of carbon dioxide testing. Carbon dioxide diffuses more readily than oxygen across gas-exchange membranes; therefore, it is assumed that a device that performs adequate oxygen transfer will also have sufficient carbon dioxide transfer. However, rigorous testing of both oxygen and carbon dioxide will be required prior to clinical translation of an oxygenator. This study is also limited by its use of adult animals in testing a device intended for premature infants. Although a premature animal model is the gold standard^8,9^, this is an extraordinarily complex animal model that requires continuous animal intensive care. For the purposes of prototyping, an adult animal model enabled rigorous testing of gas transfer in vivo. Unlike premature animals, an adult large animal enables frequent blood draws to gather a sufficiently large sample size to accurately evaluate oxygen flux. Finally, although we have incorporated design principles that have been shown to improve hemocompatibility, this study did not conduct a rigorous assessment of hemocompatibility based on consensus guidelines for the evaluation of hemocompatibility in medical devices (ISO 10993) and blood oxygenators (ISO 7199). Although this type of rigorous testing is required before determining the true hemocompatibility of our oxygenator, additional development of the blood-contacting material properties needs to be performed to maximize the hemocompatibility of the device. This includes coating the PDMS and housing with coatings such heparin^28^, polyethylene glycol^20^, nitric oxide^29^, or zwitterionic molecules^30^.

**Table 2:** Comparison of the silicon membrane oxygenator to hollow fiber membrane and flexible microfluidic oxygenators.

| Variable | Silicon membrane | Hollow-fiber membrane<sup>a</sup> | Flexible microfluidics<sup>b</sup> |
| --- | --- | --- | --- |
| Resistance (mmHg·mL<sup>-1</sup>·min) | 0.11-0.15 | 0.04 | 2.6-100 |
| Priming volume (mL) | 18 | 38 | 0.47-2.1 |
| Membrane surface area (cm<sup>2</sup>) | 57.6 | 3800 | 62.5-179 |
| O<sub>2</sub> flux (mL·min<sup>-1</sup>)<sup>c</sup> | 0.049 | 0.75 | 0.3-1 |
| Area-normalized flux (mL·min<sup>-1</sup>·m<sup>2</sup>) | 8.5 | 2.0 | 23.8-88 |
| Anticoagulation | No anticoagulation | High dose (ACT<sup>d</sup> = 200 s) | Very high dose (ACT = >250 s) |
| Flow path | Optimized flow path over entire device | Uncontrolled | Variable. Several use biomimetic branching within a single layer |

<sup>a</sup> Based on the neonatal Quadrox oxygenator <sup>35</sup>
<sup>b</sup> Based on representative devices from Potkay, Borenstein, and Selvaganapathy groups <sup>21,36–40</sup>
<sup>c</sup> Oxygen flux at a blood flow rate of 10 mL/min. Value estimated by linear extrapolation when not specifically tested at that flow rate
<sup>d</sup> Activated clotting time

In conclusion, this study represents a major step towards the development of a hemocompatible oxygenator for an artificial placenta. We show that the fundamental concept of combining a rigid flat-plate semi-permeable membrane and CFD modeling is a viable strategy to build a device that can provide efficient gas transfer while maintaining laminar blood flow, avoidance of recirculation, low pressure drop, and minimal high-shear stress.

## Methods

### Computational Fluid Dynamic and Mass Transfer Modeling

Computational fluid dynamic (CFD) modeling work was performed using Ansys software (Ansys, Canonsburg, PA, USA). The blood flow path was built in Solidworks (Dassault Systèmes, Vélizy-Villacoublay, France), imported into Ansys Meshing, and converted into a mesh containing 37.4 million elements. The inlet, outlet, and turn-around connection regions were meshed with prism layers along the surfaces and tetrahedral-shaped internal elements. The straight channel sections were meshed with hexahedral-shaped elements. Ansys CFX was used to conduct the laminar flow CFD modeling. A density of 1060 kg/m^3^ and a Cross non-Newtonian viscosity model were used to define blood properties^31,32^. Oxygen flux modeling was performed using a mass transfer model coded in Matlab (MathWorks, Natick, MA, USA). The mathematical derivation of the oxygen mass transfer model is described in detail in Dharia et al.^17^, and custom Matlab files are available for download in the Supplementary material. The modeling constants used for both the CFD and mass transfer models are summarized in Table 3 and are based on known properties of fetal blood^33^.

**Table 3:** Computational Fluid Dynamic and Mass Transfer Modeling Constants

| Variable | Value | Units |
| --- | --- | --- |
| **<u>Blood (Fetal Hemoglobin)</u>** |  |  |
| Dynamic Viscosity | $3.5 \times 10^{-3}$ | $\text{Pa} \cdot \text{s}$ |
| Density | 1060 | $\text{kg/m}^3$ |
| Oxygen Diffusivity | $1.97 \times 10^{-5}$ | $\text{cm}^2/\text{s}$ |
| Hemoglobin Concentration | 15 | $\text{g/dL}$ |
| P50 | 18.4 | $\text{mmHg}$ |
| Cooperative Binding Factor | 2.8 |  |
| **<u>PDMS</u>** |  |  |
| Density | 997 | $\text{kg/m}^3$ |
| Oxygen Diffusivity | 309.5 | Barrer |
| **<u>Testing Conditions</u>** |  |  |
| Temperature | 37 | Celsius |
| Inlet Oxygen Partial Pressure | 18 | $\text{mmHg}$ |
| Inlet Pressure (Umbilical Artery Pressure) | 20 | $\text{mmHg}$ |
| Sweep Gas Pressure | 5 | $\text{PSI}$ |
| Blood Flow Rate | 200 | $\text{mL/min}$ |

For the platelet stress damage accumulation simulation, 1000 streamlines uniformly spaced at the inlet were generated. The normalized power-law stress for each streamline was calculated via the following equation Eq. (1),^16^

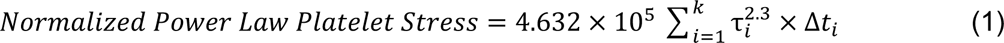

where i represents a simulated time-step, k is the total number of time-steps for a particular streamline, τ_i_ is the shear stress during that time step, Δt_i_ represents the time-step length (s). The multi-pass normalized accumulated stress was calculated by choosing a random streamline for each of 3000 passes. This process was repeated 1 million times to generate a normally-distributed probability density function demonstrating the probability of exceeding the platelet activation threshold of 1.

### Silicon membrane fabrication

Silicon on insulator (SOI) wafers were custom-designed and purchased from MEMS Engineering & Material, Inc. (Sunnyvale, CA, USA). They contained a “device” layer of 100 μm, a buried oxide layer of 1 μm, and a “handle” layer of 299 μm for an overall thickness of 400 μm. A layer of photoresist was spin-coated onto the device layer of the wafer and patterned with pores 10 μm wide by 50 μm long, separated by 10 μm on all sides. The device layer was then etched with deep reactive ion etching (DRIE) to the buried oxide layer, for an overall aspect ratio of 10:1. On the handle layer, the process was repeated except with larger 1 mm by 6 mm by 299 μm windows (width x length x depth). Following etching, the wafers were diced into 32 mm x 65 mm membranes. After dicing, the membranes were wet-etched using 49% hydrofluoric acid, which dissolved the 1 μm buried oxide layer, thereby connecting the pores with the windows and completing the semiconductor silicon membrane backbone.

To build the 5 μm layer of PDMS, first an initial sacrificial layer of approximately 250 μm PDMS (Sylgard 184, Dow Corning, Midland, MI, USA) was spin coated onto a silicon wafer (10:1 base to crosslinker mix ratio, 500 RPM for 20 seconds, cure at 80°C for 2 hours). After curing, the surface was treated with oxygen plasma (Expanded Plasma Cleaner, Harrick, Ithaca, NY, USA) at 30 Watts for 20 seconds to increase surface wettability. A second sacrificial layer of polyvinyl alcohol (PVA; Sigma-Aldrich, St. Louis, MO; powder diluted 5% w/w in water) was spin coated on top of the PDMS layer (1000 RPM for 60 seconds, cure at 60°C for 1 hour). Finally, the 5 μm layer of PDMS was created by diluting a 10:1 mixture of PDMS with an equal amount of hexane (Sigma-Aldrich, St. Louis, MO) and spin coating the PMDS/hexanes mixture onto the PVA layer (5000 RPM for 300 seconds, cure at 80°C for 2 hours).

The final composite membrane was created by combining the silicon membrane backbone and the thin PDMS layer. The PDMS construct was peeled off the silicon wafer and treated with oxygen plasma along with the silicon backbone (30 W for 20 s). The PDMS construct was then wetted with distilled water, and the 5 μm side was placed in contact with the pores side of the semiconductor silicon backbone, taking care to eliminate air bubbles. The pieces were bonded together under moderate weight on a hotplate for 12 hours at 70°C. After bonding was complete, the excess PDMS beyond the borders of the silicon chip was cut away. The composite membrane was submerged in distilled water at 70°C for 4 hours to dissolve the PVA and release the sacrificial PDMS layer, leaving the completed composite membrane.

Prior to use in the oxygenator, all membranes were individually tested for defects using a custom flow cell. Water was pumped into a chamber on the front “PDMS” side, and compressed air was pressurized on the back side up to a total of 15 PSI. The location of any bubbling was noted as a defect and covered with epoxy (Epo-Tek OD2002, Epoxy Technology, Billerica, MA, USA) on the window side. This process was repeated until the membranes were able to withstand 15 PSI with no bubbling for at least 30 seconds.

### Oxygenator Construction

To create the stack, individual composite membranes were bonded to molded gasket spacers. The gaskets were created using PDMS (Sylgard 184, 10:1 mix ratio) poured into custom-machined aluminum molds. Blood channel gaskets dimensions were 2.5 mm by 65 mm by 2 mm (width x length x height). Sweep gas gaskets were all 700 μm in height. Edge gaskets were L-shaped and 32 mm on the short arm and 57 mm on the long arm to allow a 4 mm opening on the sides for the sweep gas inlet and outlet. Additional gaskets of 3 mm width and 50 mm length were placed in the middle for structural support. The gaskets were all bonded to a single side of a chip (blood gaskets to the front side and sweep gas gaskets to the back side) using PDMS crosslinker^34^. Prior to bonding, all gaskets were measured using a micrometer, and only included if the dimensions were within 1% of the target measurement. The crosslinker was spin coated (1000 rpm for 60 seconds) on a silicon wafer to form a thin layer or approximately 3-5 μm. The gaskets were then “stamped” onto the crosslinker and aligned using a custom-machined alignment rig with the uncured crosslinker face up. The silicon membranes were lightly pressed onto the gaskets. Finally, the silicon membrane with aligned gaskets was placed onto a hotplate and cured under moderate weight (80°C for 4 hours). Once gaskets were bonded a single side, the stamping process was repeated on the other side of the gaskets to form the full stack. The entire stack was composed of 8 silicon membranes flanked by 2 silicon solids (no pores or PDMS).

Once the stack was constructed, it was placed into a 3D printed housing by sliding the stack along rails printed in the housing, which ensured proper interfacing of the blood channels in the housing and the stack. Sweep gas headers were attached to either side of the stack to connect the main sweep gas inlet and outlet of the housing to the 4 mm openings between the L-shaped gaskets in the sweep gas channels of the stack. The stack was then sealed on all edges with medical epoxy (EP30MED, Masterbond, Hackensack, NJ, USA). Two side plates and a top plate were screwed in place for additional mechanical support.

### Benchtop Water Testing

Oxygenators were initially tested for oxygen flux and pressure drop using distilled water pumped through a mock circuit loop. Although oxygen flux into blood is more complex than simple diffusion due to the red blood’s cell role as an oxygen carrier, testing in water provided value as a quality control test prior to blood testing. It enabled evaluation for leaks or gas emboli and provided a way to readily measure oxygen flux without biologically contaminating the device. A water reservoir was filled with distilled water and sparged with nitrogen for 2 minutes to achieve a partial pressure of oxygen between 50 and 100 mmHg. The water was then pumped using a peristaltic pump (Masterflex L/S Series, Cole-Parmer, Vernon Hills, IL, USA) through the device at a rate of 10 to 40 mL/min. Pressure drop was measured using two high accuracy digital pressure gauges (GE Druck DPI 104, Cole-Parmer, Vernon Hills, IL, USA) placed before and after the device. The oxygen concentration was measured in μmol/L using an optical oxygen sensing probe (NeoFox, Ocean Insight, Orlando, FL, USA). After measuring the baseline oxygen concentration, the oxygen sweep gas was turned on and pressurized to 1 PSI. The oxygen concentration was allowed to stabilize over 1 minute, and the value recorded. Oyxgen flux (J) at standard temperature and pressure (STP) was calculated using Eq. (2), where q_w_ is the volumetric flow rate of water (mL/min), R is the ideal gas constant, and n_f_ and n_i_ are the respective final and initial molar concentrations of oxygen (μmol/L) measured by the oxygen probe.

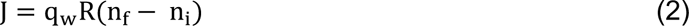

### Extracorporeal Pig Testing

Testing of the oxygenator was performed in an extracorporeal pig model at Covance Lab (San Carlos, CA, USA). All procedures were approved by the Institutional Animal Care and Use Committee at Covance. Adult Yucatan pigs were anesthetized and endotracheally intubated. Intraoperative monitoring included continuous pulse oximetry, arterial blood pressure measurements, and cardiorespiratory monitoring. A 13 French double lumen catheter (BD, Covington, GA, USA) was surgically placed into the jugular vein and sutured in place. Immediately prior to insertion of the catheter, a single dose of heparin (100 units/kg) was given to the pig. The animal was not anticoagulated through the remainder of the experiment. The extracorporeal circuit was primed with 0.9% sodium chloride solution and then connected to the venous catheter. Blood was pumped through the circuit using a Masterflex peristaltic pump at a rate of 10 and 20 mL/min. Pressure drop was meausured using pressure transducers (Edwards Lifesciences, Irvine, CA, USA) placed before and after the device. For oxygen flux measurements, 100% oxygen sweep gas was turned on and pressurized to 0.5-1 PSI. Blood samples were taken from ports before and after the device, and blood gas analysis was performed using an i-Stat blood gas analyzer (Abbott Point of Care Diagnostics, Princeton, NJ, USA). Volumetric oxygen content ([O_2_]) of the blood (mL/dL) was calculated using Eq. (3), where Hgb is the hemoglobin concentration (g/dL), S_O2_ is the oxygen saturation, and P_O2_ is the partial pressure of dissolved oxygen (mmHg). The flux was then determined by multiplying the change in oxygen content by the blood flow rate.

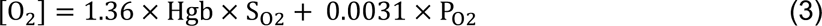

At the completion of the study, the oxygenator was gently cleared of unclotted blood by pumping normal saline at 10 mL/min. The oxygenator, which was constructed from optically clear housing material, was visually inspected for evidence of thromboses.

## Supporting information

Supplementary Files - Matlab modeling scripts

## Acknowledgements

We thank Jimmy Ly for his expert guidance in constructing and sealing the oxygenator. Dr. Blauvelt was supported in part by NIH T32 train grants (HD049303 and HL007544). The project was funded by a West Coast Consortium for Technology and Innovation grant (P50FD006425), UCSF Benioff Children’s Hospital Silver Award funds from the UCSF-Stanford Pediatric Device Consortium Annual Accelerator Competition (2020), and an NIH/NIBIB grant (U01EB025136).

## Author Contributions

DB: Concept, design, data collection, data analysis, data interpretation, drafting of the article, funding secured; NH: Data collection, data analysis, data interpretation, critical revision of the article; BD: Design, data collection, critical revision of the article; MG: CFD data analysis, CFD data interpretation, critical revision of the article; NW: Concept, design, critical revision of the article; CB: Concept, critical revision of the article. JM: Animal study protocol development, animal surgeries; BC: Membrane design, membrane fabrication, critical revision of the article; FB: Membrane fabrication, critical revision of the article; PO: Concept, critical revision of the article; SR: Concept, design, data interpretation, critical revision of the article, approval of article, funding secured.

## Data Availability

The datasets generated during the current study are available from the corresponding author on reasonable request. Matlab software codes are available for download in the Supplementary materials.

