## Supplementary Files - Matlab modeling scripts for "A Rigid Parallel-Plate Artificial Placenta Oxygenator with a Hemocompatible Blood Flow Path": FluxModel_blood.m

```
function Flux = FluxModel_blood(chip_gen,chip_size,qL,h,z_pdms,P_sweep,PP_inlet,n_membranes,chips_in_series)

% Direct questions to

%%%%%%%%%%%%%%%% EXAMPLE RUN %%%%%%%%%%%%%%%%%%%%%%%%%%%%
% To run the sample function, use the following example command:
% Flux = FluxModel_blood('MPA6', '32x65',200,2,5,5,18,2,4);


%%%%%%%%%%%%%%%% Function Input Variables %%%%%%%%%%%%%%%%
%   chip_gen: membrane generation, e.g. 'MPA6'
%   chip_size: membrane size, e.g. '32x65' 
%   qL: volumetric flow rate, in mL/min
%   h: channel height in mm
%   z_pdms: PDMS thickness in microns;
%   P_sweep: sweep gas pressure in PSI 
%   PP_inlet: partial pressure of inlet in mmHg (torr)
%   n_membranes: number of membranes per channel
%   chips_in_series: number of membranes in series


%%%%%%%%%%%%%%%%%%%%%%%%%%%%%%%%%%%%%%%%%%%%%%%%%%%%
dx = 0.001; %defines in mm, the size of each x segment. In other words, if 0.001, divides chip into 1 micron segments.
%%%%%%%%%%%%%%%% Chip Charcteristics %%%%%%%%%%%%%%%%

%Open corresponding chip file 
load([cd,'\',chip_gen,' ',chip_size,'.mat']);

%When creating new membrane files, must have the following variables:
%   w: width of the blood channel in mm. This does not include the space where the gasket blocks the blood channel
%   L: length of the blood channel in mm
%   Nwindows: number of windows
%   window_starts: numeric array with x value for the start of each large window in mm
%   window_length: length of each large window in mm
%   window_width: width of each large window in mm
%   xbar_width: space between each large window in mm along the x-axis
%   border_width: width of the space (in mm) with no membrane area along the edges of the blood channel
%   dead_area: how much of the large window is occupied by non-active membrane material (i.e. not small window area)
%   dead_area_avg_diffusion_distance: approximate average distance oxygen has to diffuse within the dead area to get to an active pore area.
%   long_chip_multiplier: number of long chips membrane space (i.e. 1x for a 10x65 chip, 3x for a 32x65 chip).


%%%%%%%%%%%%%%%% Constants %%%%%%%%%%%%%%%%

% %%% for blood model (ADULT)
% mu_blood = 0.0035; %blood dynamic viscosity in Pa*s = kg/(m·s)
% rho_blood = 1060; %blood density in kg/m^3
% hgb = 14; %hemoglobin concentration of an average adult in g/dL
% hgb_MW = 64450; %molecular weight of hemoglobin in g/mol
% hgb_MC = hgb * 10 / hgb_MW; %molar concentration of hemoglobin in moles/L
% CH_sat = hgb_MC * 4; %molar oxygen concentration (mol/L) of fully saturated hemoglobin. Each hgb molecule can hold 4 oxygen molecules
% p50_hgb = 26; %partial pressure of oxygen in mmHg at which 50% of hemoglobin molecules are bound
% n = 2.34; %hemoglobin cooperative binding factor. 
% H = 42000; %Henry's law coefficient for oxygen in water in mol H20*atm/mol O2 at 37C

%%% for blood model (FETAL)
mu_blood = 0.0035; %blood dynamic viscosity in Pa*s = kg/(m·s)
rho_blood = 1060; %blood density in kg/m^3
hgb = 15; %hemoglobin concentration of an average fetus in g/dL
hgb_MW = 64450; %molecular weight of hemoglobin in g/mol
hgb_MC = hgb * 10 / hgb_MW; %molar concentration of hemoglobin in moles/L
CH_sat = hgb_MC * 4; %molar oxygen concentration (mol/L) of fully saturated hemoglobin. Each hgb molecule can hold 4 oxygen molecules
p50_hgb = 18.4; %partial pressure of oxygen in mmHg at which 50% of hemoglobin molecules are bound
n = 2.8; %hemoglobin cooperative binding factor. 
H = 42000; %Henry's law coefficient for oxygen in water in mol H20*atm/mol O2 at 37C


%%% for water and blood models
cL = 55.5; %molar concentration of water
rho_o2 = 1.429; %oxygen density (at stp) in kg/m^3
molmass_o2 = 0.032; %molar mass O2 in kg/mol.
Dinf = 0.0000197; %oxygen bulk diffusion coefficient in cm^2/s
Pm_pdms = 309.5*10^-10; %oxygen permeability of PDMS in mL cm cm^-2 s^-1 cmHg^-1 (from Dharia et al.)
z_pdms = ((z_pdms*active_area)+((z_pdms+dead_area_avg_diffusion_distance)*dead_area))/LW_area; %accounts for diffusional distance of dead area within the large windows of the membranes


%%%%%%%%%%%%%%%%%%%%%%%%%%%%%%%%%%%%%%%%%%%
%%%%%%%%%%%%%%%%%%%%%%%%%%%%%%%%%%%%%%%%%%%

%%% Initial calculations to determine kL (liquid phase mass transfer coefficient) %%%%
%Calculate Reynold's number
dh = 4*(w/1000)*(h/1000)/(2*((w/1000)+(h/1000))); %hydraulic diameter for a rectangular channel in meters. 
dh2 = dh*(2/n_membranes); %2/n_membranes corrects for a channel lined with only one membrane (oxygen has to diffuse all the way to the other side of the channel, effectively doubling diffusion distance versus circular pipe or 2-sided channel)

A = w/1000*h/1000; %cross sectional area in m^2
v = (qL/60/100^3)/A; %velocity of water in m/s
Re = rho_blood*v*dh2/mu_blood; %Reynold's number, dimensionless

%Calculate Schmidt's number
Sc = mu_blood/(rho_blood*Dinf/100^2); %dimensionless

%Calculate Sherwood's number (dimensionless).
Sh = 1.65*(Re*Sc*dh2/(L/1000))^(1/3); %Dharia et al.

%Calculate liquid mass transfer coefficient in m/s
kL = Sh*(Dinf/100^2)/dh2; 

%Convert kL into mol s^-1 m^-2 atm^-1
kL = kL/H; %divide by Henry's constant to achieve units of m mol O2 s^-1 mol H20^-1 atm^-1
kL = kL*cL*1000; %multiply by molar concentration of water, then 1000 (L->m^3) to get to units of mol O2 s^1 m^-2 atm^-1


%%%%%%%%%%%%%%%%%%%%%%%%%%%%%%%%%%%%%%%%%%%
%%%%%%%%%%%%%%%%%%%%%%%%%%%%%%%%%%%%%%%%%%%

%Flux calculations

%divide the chip into segments
X = dx:dx:L; 

%Assign zone numbers for each X value based on the membrane characteristics
%Zone 1 = region 1,2 in Dharia et al.
%Zone 2 = region 3,4 in Dharia et al.
%Zone 3 = region 5,6 in Dharia et al.
Z=zeros(1,length(X));
Z(X<window_starts(1))=1; %All values before first window are zone 1
Z(X>(window_starts(Nwindows)+window_length))=3; %All values after last window are zone 3
for window = window_starts
    Z(X>=window & X<=(window+window_length))=2; %All windows are in zone 2
    Z(X>=window-xbar_width/2 & X<window) = 1; %Half of the bar before the window is zone 1
    Z(X>window+window_length & X<=window+window_length+xbar_width/2)=3; %Half of the bar after the window is zone 3
end

%Main loops: going through each X-segment and determining change in flux.

%Main loop #1: flux for window-containing area, i.e. regions 2, 4, and 6 
%in Dharia et al.
P_start = PP_inlet; %starting pressure =  partial pressure of inlet
wL = window_width; % for this loop, channel width (wL) is equal to the membrane window width
J_even = zeros(chips_in_series, length(X));
for c = 1:chips_in_series
    for x = 1:length(X)

        %calculate deltaM = diffusion distance through membrane in mm
        if Z(x) == 1 || Z(x) ==3 %if segment is in Zone 1 or 3

            %find distance to next nearest window
            i = window_starts-X(x);
            dist_next_window = min(i(i>0));

            %find distance to last nearest window
            i= X(x)-(window_starts+window_length);
            dist_last_window = min(i(i>0));

            %total gas diffusion (in mm) distance is PDMS thickness plus distance to nearest pore
            deltaP_M = z_pdms/1000 + min([dist_next_window,dist_last_window]);

        else %if segment is in Zone 2

            deltaP_M = z_pdms/1000;

        end

        %calculate mass transfer coefficient through PDMS membrane
        kM = Pm_pdms/(deltaP_M/10)*(100^2); % gas transfer coefficient for PDMS in mL(O2) m^-2 s^-1 cmHg^-1

        %Convert Km to units of mol s^-1 m^-2 atm^-1
        %Divide by 100^3 to convert mL->m^3, multiply by 76 to convert cmHg -> atm 
        %Then multiply by density of oxygen (in kg/m^3), divide by molar weight of oxygen (in kg/mol) to get to mol s^-1 m^-2 atm^-1
        kM = kM / 100^3 * 76 * rho_o2 / molmass_o2;

        %calculate overall mass transfer coefficient in mol s^-1 m^-2 atm^-1
        k0 = 1/(1/kM+1/kL);

        %calculate delta pressure in atm
        deltaP_t = (P_sweep*0.068+1)-P_start/760; %convert sweep gas PSI to atm and add 1 atmosphere. Sweep gas pressure is pressure on top of 1 atm. Subtract Pstart in torr after converting to atm

        %calculate flux in mol/s
        J = k0 * deltaP_t * ((wL/1000)*(dx/1000)*n_membranes); %flux = k*deltaP * surface area. Surface area = channel width*channel section length(dx)*n_membranes
        J_even(c,x) = J; %vector of flux over the chip in mol/s

        %convert flux to mmHg*L/s
        J = J * H / cL * 760; %multiply by henry's constant and divide by molar concetration of water to get atm*L/s. then convert to mmHg*L/s

        %calculate dP_Lb in mmHg, by dividing J by flow rate in L/s
        qL_channel = qL * wL/w; %the amount of flow going through this channel is equal to the ratio of the membrane channel width to the whole blood channel width
        dP_Lb = J /(qL_channel/1000/60); %qL needs to be converted from mL/min -> L/s

        %determine percentage of the partial pressure change that goes to
        %the hemoglobin sink. This piece of the loop uses Hill's equation,
        %which is the relationship betwen oxygen saturation and partial
        %pressure. Taking the Hill equation (O2sat% = P^n / p50_hgb^n + P^n)
        %and multiplying by the molar concentration of bound oxygen in 100%
        %saturated blood gives a relationship between partial pressure and
        %bound oxygen concentration. The derivative of this describes how bound
        %oxygen concentration changes as a function of changing partial
        %pressure. This reduces to the equation: dBound = (CH_sat*n*p50^n)*p^(n-1)/(p^n+p50^n)^2
        %The concentration of dissolved oxygen changes based on Henry's law:
        %[O2_unbound] = p*cL/H. The differential of this is dUnbound = cL/H.
        %The ratio of this is what we're interested in: m=dBound/dUnbound.
        pp=P_start;
        dBound = (CH_sat * n * p50_hgb^n) * pp^(n-1) / (pp^n+p50_hgb^n)^2;
        dUnbound = cL/(H*760);
        m = dBound/dUnbound; %ratio of the differential of bound versus unbound O2
        dP_Lb = dP_Lb / (1+m); %adjust change in pressure by amount bound by hemoglobin sink
        
        

        %calculate ending pressure and assign to array as well as P_start
        %for the next cycle of the loop
        P_end = dP_Lb + P_start;
        P_start = P_end;
    end
end


%main loop #2: Same as above, except calculating flux for border area, 
%i.e. regions regions 1, 3, and 5 in Dharia et al.
%Will have to double the flux at the end to account for top and bottom
%borders.
P_start = PP_inlet; %starting pressure =  partial pressure of inlet
wL = border_width; % for this loop, channel width (wL) is equal to the membrane window width
J_odd = zeros(chips_in_series, length(X));
for c = 1:chips_in_series
    for x = 1:length(X)

        %calculate deltaM = membrane thickness
        if Z(x) == 1 || Z(x) ==3 %if segment is in Zone 1 or 3 (Regions 1 and 5)

            %find distance to next nearest window
            i = window_starts-X(x);
            dist_next_window = min(i(i>0));

            %find distance to last nearest window
            i= X(x)-(window_starts+window_length);
            dist_last_window = min(i(i>0));

            %distance to nearest pore
            nearest_pore = min([dist_next_window,dist_last_window]);

            %create function to integrate
            fun = @(y) (nearest_pore^2+y.^2).^(1/2);

            %integrate function over 0 to border_width
            integ = integral(fun,0,border_width);

            %total gas diffusion distance is PDMS thickness plus
            %integral/border_width
            deltaP_M = z_pdms/1000 + integ/border_width;

        else %if segment is in Zone 2 (Region 3)

            deltaP_M = z_pdms/1000+border_width/2;

        end
        
        %calculate mass transfer coefficient through PDMS membrane
        kM = Pm_pdms/(deltaP_M/10)*(100^2); % gas transfer coefficient for PDMS in mL(O2) m^-2 s^-1 cmHg^-1

        %Convert Km to units of mol s^-1 m^-2 atm^-1
        %Divide by 100^3 to convert mL->m^3, multiply by 76 to convert cmHg -> atm 
        %Then multiply by density of oxygen (in kg/m^3), divide by molar weight of oxygen (in kg/mol) to get to mol s^-1 m^-2 atm^-1
        kM = kM / 100^3 * 76 * rho_o2 / molmass_o2;

        %calculate overall mass transfer coefficient in mol s^-1 m^-2 atm^-1
        k0 = 1/(1/kM+1/kL);

        %calculate delta pressure in atm
        deltaP_t = (P_sweep*0.068+1)-P_start/760; %convert sweep gas PSI to atm and add 1 atmosphere. Sweep gas pressure is pressure on top of 1 atm. Subtract Pstart in torr after converting to atm

        %calculate flux in mol/s
        J = k0 * deltaP_t * ((wL/1000)*(dx/1000)*n_membranes); %flux = k*deltaP * surface area. Surface area = channel width*channel sectiion length(dx)
        J_odd(c,x) = J; %vector of flux over the chip in terms of mol/s

        %convert flux to mmHg*L/s
        J = J * H / cL * 760; %multiply by henry's constant and divide by molar concetration of water to get atm*L/s. then convert to mmHg*L/s

        %calculate dP_Lb, by dividing by flow rate in L/s
        qL_channel = qL * wL/w; %the amount of flow going through this channel is equal to the ratio of the channel width to the whole blood channel width
        dP_Lb = J/(qL_channel/1000/60); %qL_channel needs to be converted from mL/min -> L/s

        %determine percentage of the partial pressure change that goes to
        %the hemoglobin sink
        pp=P_start;
        dBound = (CH_sat * n * p50_hgb^n) * pp^(n-1) / (pp^n+p50_hgb^n)^2;
        dUnbound = cL/(H*760);
        m = dBound/dUnbound; %ratio of the differential of bound versus unbound O2
        dP_Lb = dP_Lb / (1+m); %adjust change in pressure by amount bound by hemoglobin sinkF

        %calculate ending pressure and assign to array as well as P_start
        %for the next cycle of the loop
        P_end = dP_Lb + P_start;
        P_start = P_end;

    end
end

if isnan(sum(J_odd))
    J_odd = 0;
end

J_tot = sum(sum(J_odd*2+J_even,2))*long_chip_multiplier; %total flux in moles/s
Flux = J_tot * molmass_o2 / rho_o2 * 100^3 * 60; %flux in mL/min of O2 (stp)
end
```
